# A chromosome-level reference genome assembly reveals extreme loss of Y-chromosomal diversity in the Saimaa ringed seal (*Pusa saimensis*)

**DOI:** 10.64898/2026.09.11.750824

**Authors:** Emmi Olkkonen, Ari Löytynoja, Zsófia Fekete, Anastasia Konstantopoulou, Danilo F. Santoro, Claudius F. Kratochwil, Jaakko Pohjoismäki

## Abstract

Saimaa ringed seals (*Pusa saimensis*) are a highly endangered freshwater pinniped species. Due to their unique evolutionary history and intense persecution over the last century, their population size and genetic diversity are low. Although mitochondrial and genome-wide variation have been extensively investigated, the paternal evolutionary history and genetic diversity of the Y chromosome remain poorly understood due to the lack of a suitable reference sequence. In this study, we take advantage of a new chromosome-level reference genome containing a curated Y chromosome and whole-genome resequencing data from 80 individuals, and characterize Y-chromosomal diversity, compare paternal and maternal lineages, and validate genomic approaches for sex determination. Read-depth analyses confirmed the accuracy of the assembled sex chromosomes, with male individuals exhibiting the expected two-fold increase in coverage across the pseudoautosomal region of chromosome X and exclusive coverage of the Y chromosome. These coverage patterns enabled reliable molecular sex determination. Phylogenetic analyses showed that both Y chromosome and mitochondrial haplotypes of the Saimaa ringed seal form distinct monophyletic clades relative to Baltic, Lake Ladoga and Arctic ringed seals, consistent with the long evolutionary isolation of the species. Similarly, the extent of genetic diversity in the Y chromosome and the mitochondrion aligned with with this long isolation history, since relative to the diversity observed across non-Saimaa ringed seals, the contemporary Saimaa population was estimated to retain only 5% of mitochondrial diversity and approximately 1% of Y-chromosomal diversity. All together, we show that chromosome-scale reference genome assemblies that include complete sex chromosomes provide a valuable resource for conservation genomics, studies of sex-specific population history, and comparative investigations of Y chromosome evolution across pinnipeds and other mammal species.

## INTRODUCTION

The Saimaa ringed seal (*Pusa saimensis*) is an endangered freshwater pinniped endemic to Lake Saimaa in southeastern Finland (Kunnasranta et al., 2021). Genomic analyses have recently shown that the Saimaa population derives from a lineage that diverged from other extant ringed seals during the Middle Weichselian, approximately 90,000–50,000 years ago (Löytynoja et al., 2025), likely originating from one or more ice-dammed refugial populations located along the eastern margin of the Fennoscandian Ice Sheet (Mangerud et al., 2004). The present-day Lake Saimaa population therefore represents the sole surviving descendant of an ancient freshwater lineage rather than a recently isolated population derived from Baltic ringed seals after the last glaciation (Löytynoja et al., 2025; Olsen et al., 2025). Following the retreat of the Scandinavian Ice Sheet, the lineage became confined to the evolving Lake Saimaa basin, where it has persisted in geographic isolation ever since. Consistent with this deep evolutionary history and its distinct morphology and ecology, the Saimaa ringed seal has recently been elevated to full species status (Löytynoja et al., 2025; https://marinemammalscience.org/science-and-publications/list-marine-mammal-species-subspecies/).

Following severe persecution, environmental pollution and anthropogenic changes to water levels that impaired breeding habitat, the population declined to fewer than 150 individuals during the 1980s (Kunnasranta et al., 2021), before recovering through intensive conservation efforts to over 500 seals today. The prolonged isolation and small population size have affected the genetic diversity of the Saimaa ringed seal, and investigations based on microsatellites, mitochondrial DNA and, more recently, genome-wide SNP markers have shown that the population has lost a substantial proportion of both its nuclear and mitochondrial variation compared with marine ringed seals (Heino et al., 2023; Löytynoja et al., 2025; Nyman et al., 2014; Olsen et al., 2025; Sundell et al., 2023; Valtonen et al., 2014; Valtonen et al., 2012). However, the relative contributions of the initial founder event associated with colonisation of Lake Saimaa, gradual genetic erosion during postglacial isolation, and the severe anthropogenic bottleneck of the twentieth century remain debated (Heino et al., 2023). Whole-genome analyses have further revealed extensive runs of homozygosity, reflecting recent inbreeding, as well as marked genetic differentiation among the geographically isolated subpopulations within the highly fragmented lake system (Sundell et al., 2023). This subdivision has important conservation implications because each subpopulation harbours unique genetic variation that could be permanently lost through local extinctions (Löytynoja et al., 2023; Sundell et al., 2023). Consequently, understanding how genetic diversity is distributed within the lake is essential for planning translocations and other conservation measures aimed at maintaining the evolutionary potential of the species.

Mitochondrial DNA variation in the Saimaa ringed seal has been studied extensively and provides a detailed picture of the maternal population history (Heino et al., 2023; Nyman et al., 2014; Valtonen et al., 2012). Fourteen mitochondrial non-coding region haplotypes have been identified in total, although several are known only from historical material and approximately 80% of extant individuals belong to only two maternal haplotypes (Heino et al., 2023). Temporal analyses of museum specimens have demonstrated recent losses of mitochondrial diversity during the twentieth century. Nevertheless, mitochondrial DNA represents only the maternal lineage and therefore provides an incomplete view of the demographic history and genetic structure of the population.

In contrast, the paternal history of the Saimaa ringed seal has remained entirely unexplored because no Y chromosome reference sequence has previously been available. The mammalian Y chromosome is inherited strictly from father to son and, apart from the small pseudoautosomal region shared with the X chromosome (Hinch et al., 2014), does not undergo meiotic recombination. As a result, most of the Y chromosome behaves as a single haplotype, making it an ideal genomic region for reconstructing male lineages, sex-biased dispersal and historical breeding patterns (Hughes & Page, 2015). At the same time, its highly repetitive structure and abundance of duplicated sequences have made the Y chromosome one of the most difficult parts of mammalian genomes to assemble (Rhie et al., 2023), leaving it absent or highly fragmented in many reference genomes.

We have previously published a chromosome-level reference genome for the Saimaa ringed seal (Grethlein et al., 2026), which includes a curated Y chromosome assembly (available through NCBI under the accession GCA_059488705.1), enabling the first investigation of paternal genetic variation in this species. The two sex chromosomes of the reference genome were originally merged by the assembler at the homologous pseudoautosomal region but were manually separated during curation of the final assembly. As it was impossible to separate the X-and Y-chromosomal haplotypes, the pseudoautosomal region was left as a part of the X chromosome. Using whole-genome sequencing data from 80 contemporary individuals, we characterise Y chromosome diversity within Lake Saimaa, compare paternal and maternal patterns of genetic variation, and demonstrate the utility of the new reference genome for molecular sex determination. Together, these genomic resources provide new insights into the evolutionary history and conservation genetics of one of the world’s most endangered pinnipeds. More generally, our chromosome-level Saimaa ringed seal reference genome, incorporating complete sex chromosomes, provides an additional genomic resource for comparative genomic studies in mammals, including Y chromosome evolution.

## MATERIALS AND METHODS

### Samples and sequencing

Samples representing the Saimaa ringed seal population across Lake Saimaa were collected from the principal water basins and subpopulations, including Pihlajavesi, Kolovesi, Haukivesi, South Saimaa, and North Saimaa. The samples comprised individuals of both sexes and covered multiple collection years, providing representation of the spatial and temporal variation within the population. Sample metadata, including locality, sex, and year of collection, are provided in Supplementary Information Table S1.

Acquisition, processing and sequencing of samples from 105 Saimaa ringed seals, 9 Ladoga ringed seals, 12 Baltic ringed seals and 25 Arctic ringed seals from Svalbard and Greenland were described by Löytynoja et al. (2023) and Rosing-Asvid et al. (2023). Out of the 151 samples, only 62 Saimaa ringed seals and 18 ringed seals from other populations were included in the analyses due to many of the individuals having low sequencing depth and/or high missingness in Y chromosome variants. Since the publication of the articles by Löytynoja et al. and Rosing-Asvid et al., out of the 80 seals included in our analyses, 57 of the Saimaa ringed seals, 2 Ladoga ringed seals, 4 Baltic ringed seals, and 1 Arctic ringed seals from Svalbard have been sequenced to higher depth at the DNA Sequencing and Genomics Core Unit at the University of Helsinki using the AVITI platform. Similarly, the 7 Arctic ringed seal samples from Greenland were further sequenced by BGI Hong Kong using the DBNSEQ machine (See “Acknowledgements”).

### WGS data processing

Mapping of the whole-genome sequencing data was carried out using a nextflow (v. 22.10.1) pipeline (https://github.com/emmiolkkonen/mapping_nextflow). Within the pipeline, alignment of short reads against the reference genome (NCBI accession JBXAED000000000) was carried out with BWA-MEM2 (Li, 2013) and samtools (1.16.1) (Danecek et al., 2021). Realignment of indels was done with GATK (3.8-1-0) (McKenna et al., 2010). Variant calling of haploid mitochondrial and Y chromosome genotypes is described below in the section “Maternal and paternal lineages: Analysis of variation in the Y chromosome and mitochondria”.

### Depth analysis

To compare sex chromosome coverage between male and female seals and assess it as a tool for sex determination, sequencing coverage was calculated in 10 MB windows, using samtools depth, for the X and Y chromosomes. 62 Saimaa ringed seals with sequencing coverage in chromosome 1 exceeding 10x were included in the depth analysis. As repetitive sequences may affect coverage estimates, we ran RepeatMasker (v.4.1.5, https://www.repeatmasker.org/RepeatMasker/) using the rmblastn search engine (v.2.14.0) and Dfam (v.3.7). (2023-01-11) database with “Canis lupus familiaris” as the query species and only included regions not covered by repeats. In addition to the sex chromosomes, we carried out the same calculation for autosomes 14 and 15. Chromosome 15 was used to normalise the depth estimates, and chromosome 14 was included as a control to assess the coverage patterns in non-sex chromosomes. Out of the 62 Saimaa ringed seals, we chose 15 for visualisation of depth across chromosomes. Out of these samples, 5 were female and 5 were male. The remaining 5 seals with unverified sex were used to test the method’s performance. The normalised windowed depths were visualised in R using ggplot2 (v. 4.0.3, Wickham, 2016).

To assess the overall distributions of X and Y chromosome coverage in males and females, we averaged the normalized depths of chromosome X, Y and 14 for the larger set of 62 Saimaa ringed seals, including 25 females and 37 males. The pseudoautosomal region at the end of the X chromosome, determined from the low pairwise differences in normalized coverages between two female and two male individuals, was excluded. The region spanned approximately 6.8 MB and is close to the size of the pseudoautosomal region in dog (Young et al. 2008).

For the Y chromosome, only scaffolds SUPER_Y and SUPER_Y_unloc_1 were included in this analysis due to unusually high depths in shorter Y scaffolds, indicative of repetitive regions or assembly errors. The normalised depths were averaged and plotted with ggplot2.

### Maternal and paternal lineages: Analysis of variation in the Y chromosome and mitochondria

We inferred phylogenies separately for the maternally and paternally inherited parts of the genome for a set of 28 male Saimaa ringed seals and 18 male ringed seals from Lake Ladoga, the Baltic Sea, and the Arctic.

In the initial mapping and variant calling of the mitochondrion, 105 Saimaa ringed seals and 46 ringed seals from other populations were included. For the Y chromosome, only samples with sequencing coverage over 10x were used, including 62 Saimaa ringed seals and 42 ringed seals from Lake Ladoga, the Baltic Sea, and the Arctic. Reads mapping to the mitochondrial regions (including erroneous partial copies in the nuclear part) were extracted from the BAM alignments, downsampled to expected coverage of 900X, and mapped back to the mitochondrial genome using BWA-MEM2. Variant calling was done with HaplotypeCaller, using parameter “-ploidy 1”, CombineGVCFs and GenotypeGVCFs from GATK 4 (https://github.com/broadinstitute/gatk). For mapping and variant calling, the same steps were carried out for the Y-chromosomal regions, except for the downsampling and remapping.

Initial quality filtering for variants was carried out as recommended by GATK (https://gatk.broadinstitute.org/hc/en-us/articles/360035890471-Hard-filtering-germline-short-variants). After mapping and variant calling, only 46 male seals with high data quality were included for the analyses.

Since the mitochondrial DNA is not expected to harbor repetitive transposable elements, biallelic SNPs within the entire mitochondrion were included. In contrast, the Y chromosome is highly repetitive, contains pseudoautosomal regions leading to erroneous read mapping, and as a haploid chromosome, has generally the lowest sequencing coverage. We used various methods to remove the potential error from those. As described by Olkkonen and Löytynoja (2023), SNPable (Li et al., 2013) was used to identify callable regions in the scaffolds SUPER_Y and SUPER_Y_unloc_1, and repeat regions (described in “Depth analysis”) were then subtracted from these unique regions, creating a positive mask of callable non-repetitive regions. Biallelic SNPs within positive masked regions were extracted, and individual- and site-level missingness was calculated using VCFtools (v. 0.1.17, Danecek et al., 2011) “--missing-indv” and “--missing-site”, and male seals and SNPs with less than 10% missingness were retained. To remove error from potentially misassembled or pseudoautosomal regions, sites with coverage above zero in a set of five females were excluded. Additionally, any variant with a non-reference call showing more than 10% of total allele depth consisting of reference read support in more than 50% of individuals was deemed impossible on a haploid chromosome and discarded. To further exclude spurious calls, alternative calls with support from less than 3 reads were considered as missing data, and sites with depth more than 1.5x the median coverage across sites were discarded. The remaining missing genotypes were assigned as the reference allele; for comparison, the phylogenetic analysis was performed by retaining the missing genotypes as “N” (Figs. S2–3). Since one of the Saimaa ringed seal samples had missing genotypes for 43% of the few variable sites within the population, this sample was excluded. The same samples were used for phylogenetic inference of the mitochondrion and Y chromosome, retaining 46 males, and 600 and 404 SNPs after filtering, respectively. The VCFs were transformed to FASTA format following Löytynoja et al. (2025), creating a multiple sequence alignment of variable positions.

RAxML Next Generation (v. 2.0.2, Kozlov et al., 2019) was used to infer phylogenetic trees for the mitochondrion and Y chromosome. First, Maximum-likelihood (ML) trees were estimated using the GTR+G model. Since only variable sites were given as input, ascertainment correction was done to account for invariable sites in the analysis. For the mitochondrion, the correction was added to the model parameter as ASC_STAM{n(A)/n(C)/n(G)/n(T)}”, incorporating the counts of each base within invariable sites, and for the Y chromosome with “+ASC_LEWIS.

The credibility of the ML trees was estimated by bootstrapping, creating 1000 bootstrapping trees for the mitochondrion and the Y chromosome. The branch support values were then mapped to the ML trees. All these steps were carried out by using the parameter “--all” in RAxML-ng. For running the three steps separately, “--msa”, “--bootstrap” and “support” can be used. The ML trees with support values were visualised with package ggtree (v. 4.3.0, Yu et al., 2017), and the cophylogeny tree using phytools (v. 2.5-2) in R. For the cophylogeny tree, iTOL (https://itol.embl.de/) was used for manually midpoint-rooting both trees. Mitochondrial and Y-chromosomal diversities – the mean pairwise genetic distances – within Saimaa relative to that across all non-Saimaa ringed seal populations was calculated using a custom R function, applying the package ape (v. 5.8-1, Paradis and Schliep, 2019).

### Code availability

The code used for the analyses described above, along with the Y chromosome gene annotations, will be available in “https://github.com/emmiolkkonen/AZF_Saimaa_analyses”.

## RESULTS

### Validating the sex-chromosome assembly of the Saimaa ringed seal reference genome

To validate the assembly of the sex chromosomes, we aligned whole-genome short-read sequencing data from individuals of known sex to the reference genome and examined sequencing coverage across a control autosomal chromosome and the X and Y chromosomes (Fig. 1). As expected for an autosomal control, chromosome 14 displayed uniform coverage in both sexes (Fig. 1a). Male individuals exhibited approximately two-fold higher read coverage over the pseudoautosomal region (PAR) of the X chromosome than over the X-specific region, reflecting the presence of homologous PAR sequences shared between the X and Y chromosomes (Fig. 1b). In contrast, females showed uniform coverage across the X chromosome, with no increase over the PAR. Although a few lower-quality samples displayed local fluctuations in sequencing coverage, the characteristic two-fold increase over the PAR was consistently observed in all male individuals. Similar coverage irregularities were also evident across the autosomal control chromosome, indicating that they are not specific to the sex chromosomes or the genome assembly. Rather, they most likely reflect post-mortem DNA degradation and the consequent uneven representation of genomic regions in sequencing libraries. Differential fragmentation and loss of DNA during degradation and library preparation can preferentially reduce coverage in some genomic regions, producing the alternating peaks and troughs observed in lower-quality samples without affecting the overall interpretation of chromosome-wide coverage patterns (Ross et al., 2013; Poptsova et al., 2014).

**Fig. 1.**
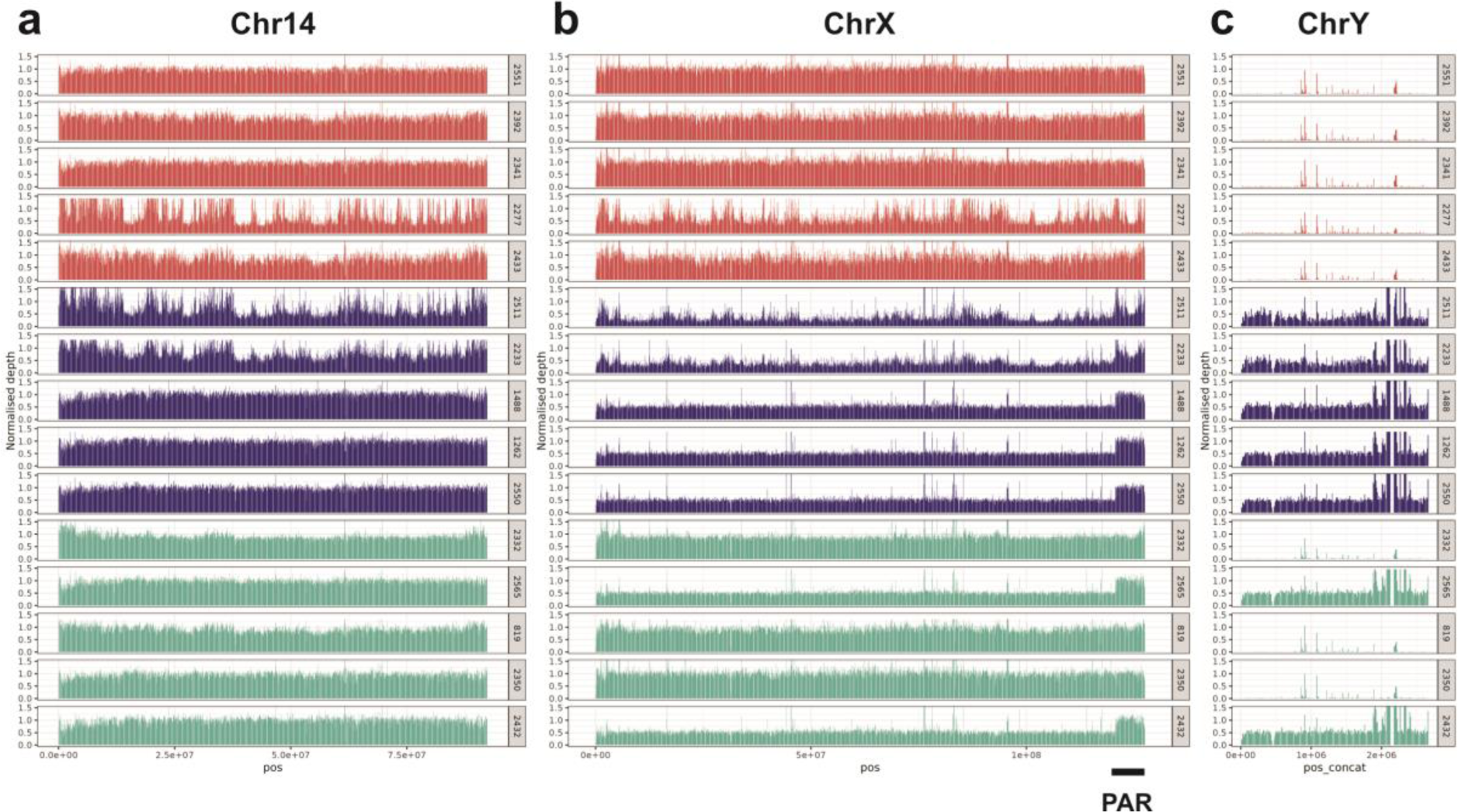
Validation of the sex chromosome assembly using whole-genome sequencing read coverage from individuals of known and unknown sex. Short-read coverage is shown across **(a)** autosomal chromosome 14 (control), **(b)** chromosome X, and **(c)** chromosome Y for four females (red; top), four males (blue; middle), and four individuals of unknown sex (green; bottom). The pseudoautosomal region (PAR) on chromosome X is indicated. Male individuals exhibit approximately two-fold higher coverage across the PAR than across the X-specific region, reflecting the presence of homologous PAR sequences on both the X and Y chromosomes. In contrast, female individuals show no increase in coverage over the PAR and virtually no read alignment to the Y chromosome. Uniform coverage across chromosome 14 in both sexes confirms that these patterns are specific to the sex chromosomes.

Coverage over the Y chromosome further distinguished the sexes. Male individuals showed uniform alignment across the assembled Y chromosome scaffolds, whereas female samples exhibited virtually no Y chromosome coverage (Fig. 1c). These coverage patterns enabled unambiguous sex assignment of specimens with unverified sex (Fig. 1, green diagrams), with the ratio of average Y-to X-chromosome coverage providing complete separation between male and female individuals (Fig. 2). As expected, the females also show substantially higher missingness across the Y-chromosomal variants (Supplementary Information Fig. S1).

**Fig. 2.**
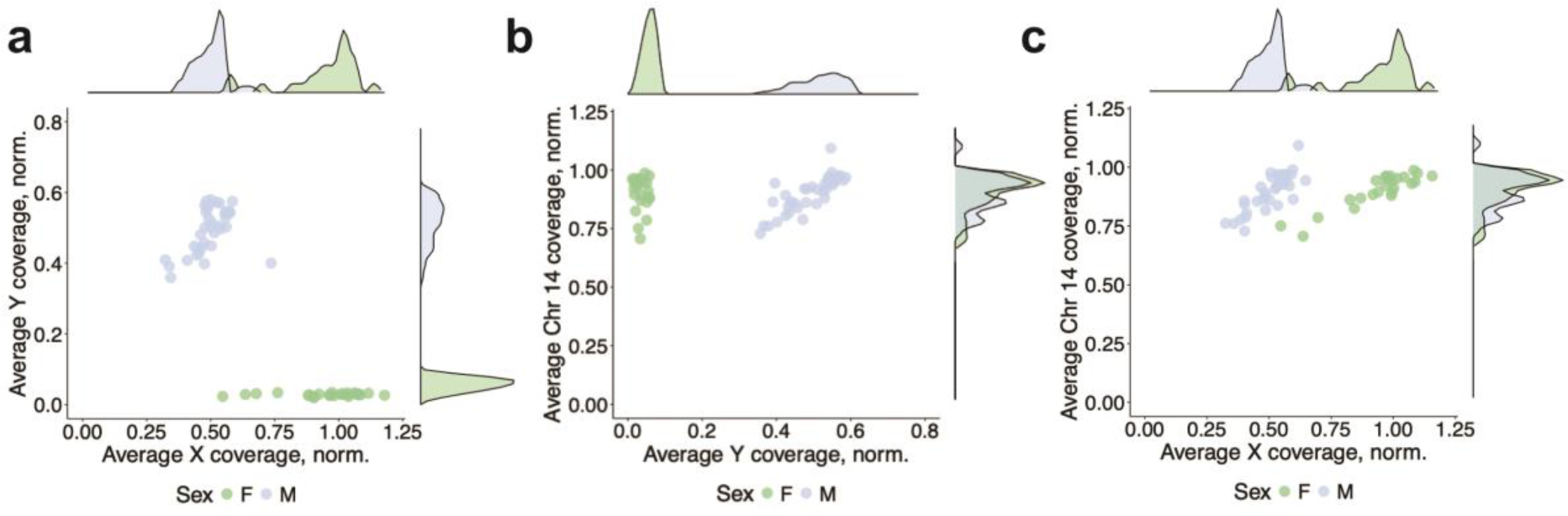
Sex assignment based on normalized sequencing coverage of the sex chromosomes. Scatter plots show the average normalized sequencing coverage of (**a**) chromosome X versus chromosome Y, (**b**) chromosome 14 (autosomal control) versus chromosome Y, and (**c**) chromosome 14 versus chromosome X for all sequenced individuals. Male (M) and female (F) specimens form two clearly separated clusters in panels **a** and **b** due to the presence or absence of Y chromosome coverage. In panel **c** males have two copies of chromosome 14 but have half of the female X-chromosome copy number. Variation in DNA quality among samples causes differences in overall sequencing coverage (see Fig. 1), leading to partial overlap between some female and male individuals.

To further validate the Y chromosome assembly, we examined its gene content. The assembly contains the expected complement of conserved mammalian Y-linked genes involved in male sex determination, spermatogenesis and testicular function (https://github.com/emmiolkkonen/AZF_Saimaa_analyses), supporting the completeness and correct assignment of the assembled Y chromosome sequence.

### Y-chromosomal diversity in Saimaa ringed seal

We next characterised Y chromosome diversity using whole-genome sequencing data from 28 Saimaa ringed seals, 5 Baltic ringed seals, 3 Lake Ladoga ringed seals and 10 Arctic ringed seals, and compared paternal lineage diversity with maternal lineage mitochondrial DNA variation in the same individuals.

As observed for mitochondrial DNA, all Y chromosomes from the Saimaa ringed seal formed a well-supported monophyletic clade that was clearly distinct from those of the Baltic, Ladoga and Arctic populations (Fig. 3, Supplementary Information Fig. S2), consistent with the long evolutionary isolation of the Saimaa lineage. In our sampling, after collapsing haplotypes represented by single individuals to their ancestral haplotype groups to account for potential noise in the low-coverage data, four Y chromosome haplotypes (arbitrarily designated A–D; Fig. 3, Supplementary Information Fig. S2 vs. S3) and five mitochondrial haplotypes (Fig. 3, Supplementary Information Fig. S4) could be identified. A total of 14 mitochondrial haplotypes are known from Saimaa, designated H1–H14, with H1 and H3 accounting for almost 80% of individuals, while some known only from historical samples (Heino et al. 2023). Traditionally, these haplotypes have been defined based on variation within the mitochondrial non-coding region rather than complete mitochondrial genome sequences. Consequently, the non-coding region haplotypes do not necessarily reflect the whole-mitogenome phylogeny. For example, H1 is being represented by two distinct lineages in our phylogeny (Fig. 3). While some geographical pattern is apparent in the distribution of both Y chromosome and mtDNA haplotypes (Fig. 4), the limited number of samples and uneven sampling across Lake Saimaa do not allow making conclusions about population structure among paternal or maternal lineages.

**Fig. 3.**
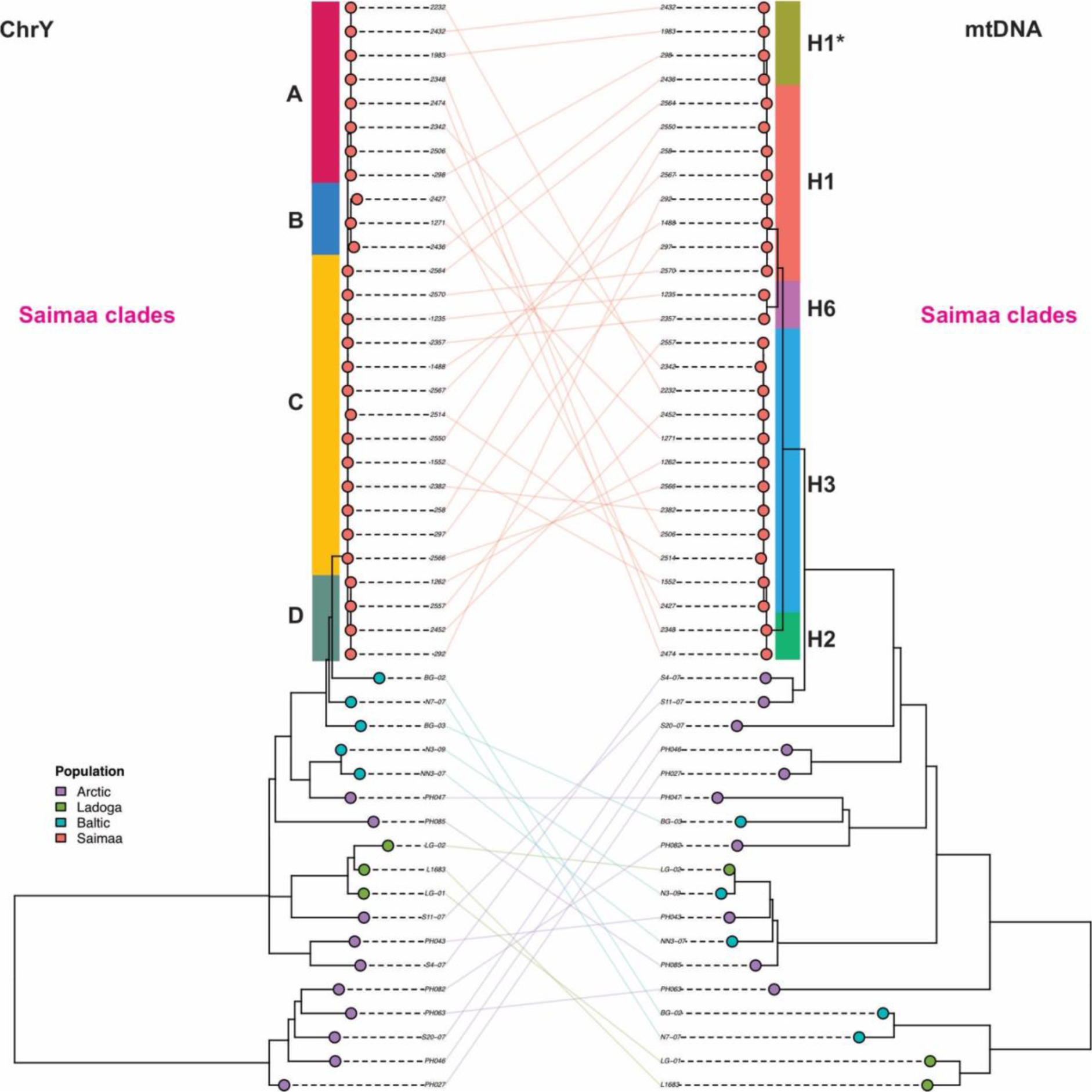
Cophylogeny of Y chromosome (ChrY; left) and mitochondrial genome (mtDNA; right) from Arctic, Baltic, Lake Ladoga, and Saimaa ringed seals. Saimaa ringed seals form distinct monophyletic clades in both the Y chromosome and mitochondrial phylogenies, reflecting their long-term evolutionary isolation. In contrast, paternal and maternal lineages from the Arctic, Baltic, and Lake Ladoga populations are intermingled, consistent with their more recent shared evolutionary history and gene flow. Lines connecting the two trees link the corresponding Y chromosome and mitochondrial genomes from the same individual. Different genetic lineages within the Saimaa population are indicated by differently coloured boxes. The ChrY haplotypes are labeled arbitrarily A–D, while the mitochondrial haplotype labels follow the established NCR haplotype nomenclature. Note that the H1 haplotype is represented by two lineages based on differences in the mitochondrial coding region. Due to the potential noise in low-coverage ChrY data, three sequence haplotypes present in single individuals were collapsed to their ancestral haplotype group.

**Fig. 4.**
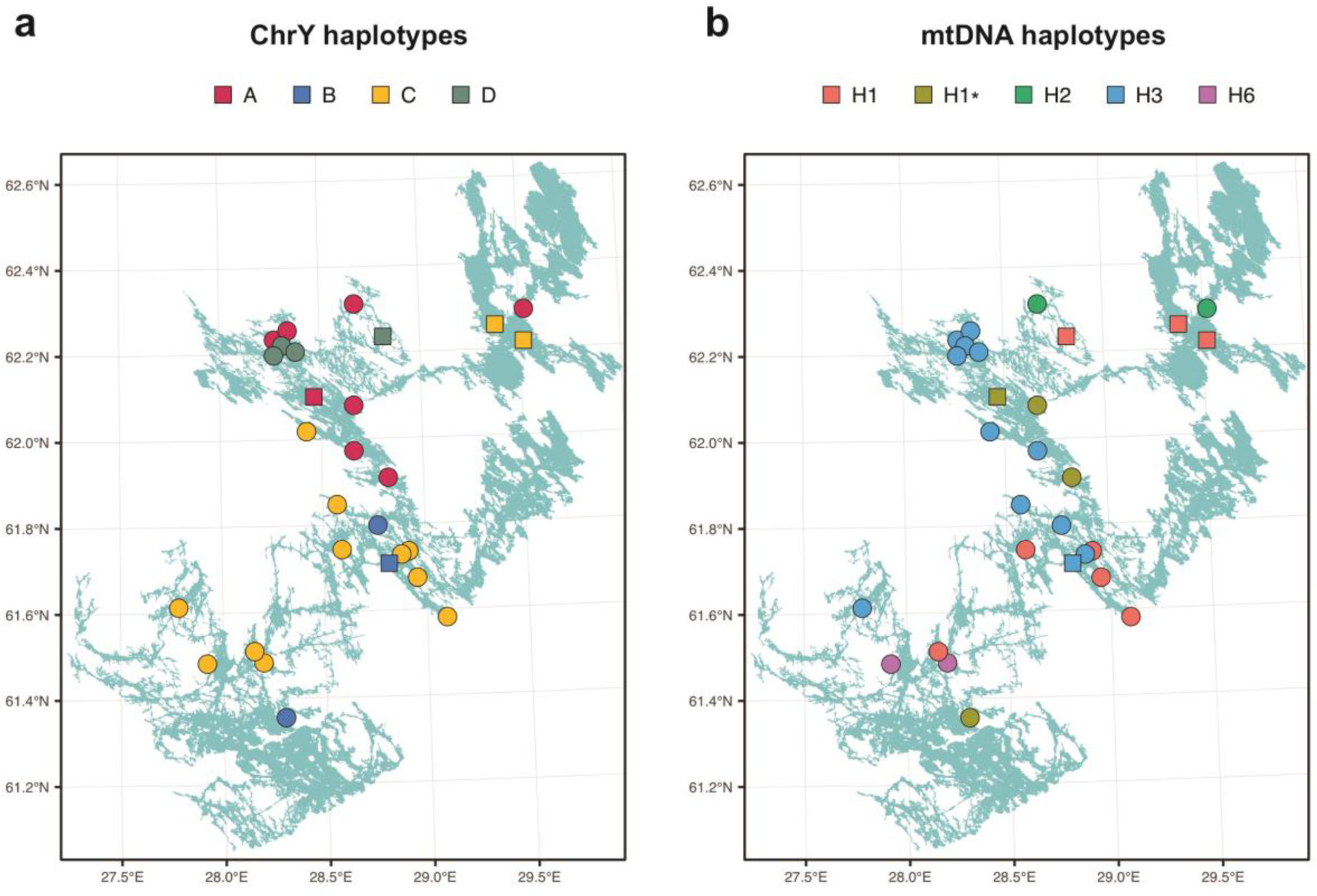
Distribution of Y chromosome (ChrY; a) and mitochondrial DNA (mtDNA; b) haplotypes among the male samples included in this study. Haplotypes are indicated by colour, as shown in Fig. 3. Individuals lacking precise sampling coordinates are indicated by squares.

To compare paternal and maternal diversity on an equivalent basis, we calculated the mean pairwise genetic distances among Saimaa individuals using both the complete mitochondrial genomes and Y chromosome sequences. Relative to the corresponding diversity observed across all non-Saimaa ringed seal populations included in the analysis, the Saimaa ringed seal retained 5% of mitochondrial diversity and 1% of Y chromosome diversity. Because both phylogenies were constructed from the same set of individuals, this comparison is not confounded by differences in sampling.

## DISCUSSION

### Sex chromosome assembly of Saimaa ringed seal

Our previously published chromosome-level reference genome includes the first Y chromosome assembly for the Saimaa ringed seal (Grethlein et al., 2026), enabling genomic analyses of paternal lineages that have previously been inaccessible. The assembly is well supported by read-depth analyses from individuals of known sex (Fig. 1, Fig. 2). As expected for a male heterogametic species, males displayed approximately two-fold higher sequencing coverage across the pseudoautosomal region of the X chromosome than across the X-specific region, reflecting homologous sequences present on both sex chromosomes. Conversely, female individuals showed essentially no coverage over the assembled Y chromosome, while autosomal coverage remained comparable between the sexes. These predictable coverage patterns demonstrate that our assembly enables reliable sex determination of individuals lacking field records and can also identify discrepancies in recorded sex assignments (Supplementary Information Fig. S1).

The assembly also contains the expected complement of conserved mammalian Y-linked genes, supporting the assignment of the principal Y scaffolds. Nevertheless, a small number of assembled Y contigs remain less well supported (Fig. 1b). Given the highly repetitive nature of mammalian Y chromosomes and the abundance of duplicated sequences shared with autosomes and the X chromosome, some contigs may represent repetitive or paralogous regions that are difficult to assign unambiguously. Future refinements based on additional long-read sequencing or comparative assemblies from related pinniped species may help resolve these regions further. Despite these uncertainties, the current assembly clearly captures the vast majority of the male-specific region of the Y chromosome and provides a robust resource for evolutionary and conservation genomic studies.

### Y chromosome diversity and male lineages of the Saimaa ringed seal

The Y chromosome phylogeny mirrors previous observations based on mitochondrial and nuclear genomes in demonstrating that the Saimaa ringed seal forms a genetically distinct lineage from Baltic, Ladoga and Arctic ringed seals (Fig. 3). The most striking finding, however, is the contrast between paternal and maternal diversity retained within Lake Saimaa. Relative to the diversity observed across the different ringed seals, the Saimaa population retains approximately 5% of mitochondrial diversity but only about 1% of Y-chromosomal diversity. Because these estimates were calculated from the same individuals, they are directly comparable and cannot be attributed to differences in sampling. Although the exact values depend to some extent on phylogenetic reconstruction, the qualitative difference is substantial and robust.

The observation is noteworthy also because the mammalian Y chromosome contains a substantially larger amount of sequence compared to the mitochondrial genome, majority of which is non-coding and therefore under weak functional constraint. Such sequences provide greater opportunity for neutral variation to accumulate (Hinch et al., 2014; Rhie et al., 2023; Tomaszkiewicz et al., 2017). Consequently, Y chromosomes generally harbour more segregating variants than mitochondrial genomes over comparable evolutionary timescales.

However, the evolvable sequence length does not determine standing diversity: the effective population size of the Y chromosome is strongly influenced by the number and reproductive success of males and is therefore substantially reduced in many mammalian species due to its uneven distribution among males (e.g. Poznik et al., 2016; Wutke et al. 2018).

Several, not mutually exclusive, demographic processes could contribute to the extent of diversity loss in the Y chromosome. First, the known male-biased dispersal (Niemi et al.,, 2013; Valtonen et al., 2012) may have influenced the spatial distribution and persistence of paternal lineages between the water basins of Lake Saimaa, while simultaneously exposing dispersing males to e.g. increased bycatch mortality, if movements bring them into areas with greater fishing pressure. Second, although the mating system of ringed seals is thought to be relatively evenly distributed among males compared with highly polygynous pinnipeds (Atkinson, 1997; Le Boeuf, 1991), variance in male reproductive success can be sufficient to reduce the effective population size of the Y chromosome and increase the probability of a lineage loss. Under a severe and prolonged population bottleneck, as experienced by the Saimaa ringed seal (Kunnasranta et al., 2021, Heino et al., 2023), combination of sex-specific differences in survival and reproduction could result in substantial stochastic loss of Y chromosome lineages even when mitochondrial lineages have been persisting.

However, the extent of paternal lineage diversity inferred here should be interpreted with caution. Although the Y chromosome is haploid and is expected to evolve in a tree-like manner, inferring phylogenetic trees from whole-genome data is a technical challenge. The first source of noise are the pseudoautosomal and repeat regions that gather erroneously mapped reads and create false variants, and the second are the stochastic errors in the variant calling process due to the low sequencing depth of the haploid chromosome. Whereas the average sequencing coverage for the mtDNA was capped around 900X, for some individuals, the mean coverage for the Y chromosome was as low as 3X based on a VCF file (mean of 3.30X and 4.60X for *P. saimensis* and *P. hispida*, respectively). Consequently, the estimated proportion of Y chromosome variation retained in Saimaa should be regarded as an approximate rather than precise quantitative measure. Importantly, however, the identification of multiple distinct Y chromosome lineages within the Saimaa ringed seal appears robust.

Although these hypotheses remain speculative, the discrepancy between mitochondrial and Y chromosome diversity demonstrates that maternal markers alone provide an incomplete picture of the evolutionary history of the Saimaa ringed seal. The availability of chromosome-scale Y chromosome data therefore opens new opportunities to investigate sex-specific demographic history, dispersal and effective population size, both in the Saimaa ringed seal and more broadly across pinnipeds.

## ACKNOWLEDGEMENTS

We thank Miina Auttila, Riikka Alakoski, Mikko Suonio and the rest of the Saimaa Ringed Seal team from Metsähallitus Parks & Wildlife Finland for providing samples throughout the years, as well as their efforts in practical conservation of this unique endemic mammal. We are indebted to Mervi Kunnasranta and Marja Niemi from the Saimaa ringed seal research group at the University of Eastern Finland for the many useful discussions and kind suggestions they made to improve our work. We also thank Paolo Momigliano (Tuscia University, Viterbo, Italy & University of Hong Kong, Hong Kong SAR, China) for providing us with deeper WGS data for Arctic ringed seals from Greenland (samples originally from Morten Tange Olsen, Globe Institute, University of Copenhagen, Denmark). Research has received funding from the European Union’s LIFE programme and projects Our Saimaa Seal LIFE (LIFE19 NAT/FI/000832) and Support of Survival LIFE (101292872 LIFE25-NAT-FI-LIFE SOS). The material reflects the views of the authors, neither the European Commission nor the CINEA is responsible for any use that may be made of the information it contains.

## AUTHOR CONTRIBUTIONS

JP conceived the study; JP, CFK supervised the study; JP, CFK provided funding; JP, CFK, AL provided materials; EO, AL, AK, ZF, DFS performed the analyses; JP, AL, CFK and EO drafted the manuscript; all authors provided editorial input and approved the final version of the manuscript.

## REFERENCES

Atkinson S. 1997: Reproductive biology of seals. --- Reviews of Reproduction 2: 175–194, doi:10.1530/ror.0.0020175.

Danecek P., Auton A., Abecasis G., Albers C.A., Banks E., DePristo M.A., Handsaker R.E., Lunter G., Marth G.T., Sherry S.T., McVean G., Durbin R. & 1000 Genomes Project Analysis Group. 2011: The variant call format and VCFtools. --- Bioinformatics 27: 2156–2158, doi:10.1093/bioinformatics/btr330.

Danecek P., Bonfield J.K., Liddle J., Marshall J., Ohan V., Pollard M.O., Whitwham A., Keane T., Davies R.M., Li H. & Durbin R. 2021: Twelve years of SAMtools and BCFtools. --- GigaScience 10: giab008, doi:10.1093/gigascience/giab008.

Grethlein M., Fekete Z., Goffart S., Kiebler A., Kunnasranta M., Niemi M., Santoro D.F., Wehrenberg G., Winter S., Prost S. & Pohjoismäki J. 2026: Chromosome-level reference genome assembly of the Saimaa ringed seal (Pusa saimensis) – an ancient glacial relict landlocked pinniped. --- bioRxiv 2026.08.13.744633, doi:10.64898/2026.08.13.744633.

Heino M.T., Nyman T., Palo J.U., Harmoinen J., Valtonen M., Pilot M., Översti S., Salmela E., Kunnasranta M., Väinölä R., Hoelzel A.R. & Aspi J. 2023: Museum specimens of a landlocked pinniped reveal recent loss of genetic diversity and unexpected population connections. --- Ecology and Evolution 13: e9720, doi:10.1002/ece3.9720.

Hinch A.G., Altemose N., Noor N., Donnelly P. & Myers S.R. 2014: Recombination in the human pseudoautosomal region PAR1. --- PLoS Genetics 10: e1004503, doi:10.1371/journal.pgen.1004503.

Hughes J.F. & Page D.C. 2015: The biology and evolution of mammalian Y chromosomes. -- Annual Review of Genetics49: 507--527, doi:10.1146/annurev-genet-112414-055311.

Kozlov A.M., Darriba D., Flouri T., Morel B. & Stamatakis A. 2019: RAxML-NG: a fast, scalable and user-friendly tool for maximum likelihood phylogenetic inference. -- Bioinformatics 35: 4453–4455, doi:10.1093/bioinformatics/btz305.

Kunnasranta M., Niemi M., Auttila M., Valtonen M., Kammonen J. & Nyman T. 2021: Sealed in a lake – Biology and conservation of the endangered Saimaa ringed seal: A review. --- Biological Conservation 253: 108908, doi:10.1016/j.biocon.2020.108908.

Le Boeuf B.J. 1991: Pinniped mating systems on land, ice and in the water: Emphasis on the Phocidae. --- In: Renouf D. (ed.), The Behaviour of Pinnipeds: 45--65. Springer Netherlands, Dordrecht.

Li H. 2013: Aligning sequence reads, clone sequences and assembly contigs with BWA-MEM. --- arXiv 1303.3997, doi:10.48550/arXiv.1303.3997.

Löytynoja A., Pohjoismäki J., Valtonen M., Laakkonen J., Morita W., Kunnasranta M., Väinölä R., Olsen M.T., Auvinen P. & Jernvall J. 2025: Deep origins, distinct adaptations, and species-level status indicated for a glacial relict seal. --- Proceedings of the National Academy of Sciences of the United States of America 122: e2503368122, doi:10.1073/pnas.2503368122.

Löytynoja A., Rastas P., Valtonen M., Kammonen J., Holm L., Olsen M.T., Paulin L., Jernvall J. & Auvinen P. 2023: Fragmented habitat compensates for the adverse effects of genetic bottleneck. --- Current Biology 33: 1009–1018.e7, doi:10.1016/j.cub.2023.01.040.

Mangerud J., Jakobsson M., Alexanderson H., Astakhov V., Clarke G.K.C., Henriksen M., Hjort C., Krinner G., Lunkka J.-P., Möller P., Murray A., Nikolskaya O., Saarnisto M. & Svendsen J.I. 2004: Ice-dammed lakes and rerouting of the drainage of northern Eurasia during the Last Glaciation. --- Quaternary Science Reviews 23: 1313–1332, doi:10.1016/j.quascirev.2003.12.009.

Niemi M., Auttila M., Viljanen M. & Kunnasranta M. 2013: Home range, survival, and dispersal of endangered Saimaa ringed seal pups: Implications for conservation. --- Marine Mammal Science 29: 1–13, doi:10.1111/j.1748-7692.2011.00521.x.

Nyman T., Valtonen M., Aspi J., Ruokonen M., Kunnasranta M. & Palo J.U. 2014: Demographic histories and genetic diversities of Fennoscandian marine and landlocked ringed seal subspecies. --- Ecology and Evolution 4: 3420–3434, doi:10.1002/ece3.1193.

Olkkonen E. & Löytynoja A. 2023: Analysis of population structure and genetic diversity in low-variance Saimaa ringed seals using low-coverage whole-genome sequence data. --- STAR Protocols 4: 102567, doi:10.1016/j.xpro.2023.102567.

Olsen M.T., Löytynoja A., Valtonen M., Knudsen S.W., Bang S., Gunnersen C., Rosing-Asvid A., Ferguson S.H., Dietz R., Kovacs K.M., Lydersen C., Jernvall J., Auvinen P. & Galatius A. 2025: Complex origins and history of the relict Fennoscandian ringed seals. -- Ecology and Evolution 15: e71067, doi:10.1002/ece3.71067.

Paradis E. & Schliep K. 2019: ape 5.0: an environment for modern phylogenetics and evolutionary analyses in R. --- Bioinformatics 35: 526–528, doi:10.1093/bioinformatics/bty633.

Poptsova M., Il’icheva I., Nechipurenko D., Panchenko L., Khodikov M., Oparina N., Polozov R., Nechipurenko Y. & Grokhovsky S. 2014: Non-random DNA fragmentation in next-generation sequencing. --- Scientific Reports 4: 4532, doi:10.1038/srep04532.

Poznik G.D., Xue Y., Mendez F.L., Willems T.F., Massaia A., Wilson Sayres M.A., Ayub Q., McCarthy S.A., Narechania A., Kashin S., Chen Y., Banerjee R., Rodriguez-Flores J.L., Cerezo M., Shao H., Gymrek M., Malhotra A., Louzada S., Desalle R., Ritchie G.R.S., Cerveira E., Fitzgerald T.W., Garrison E., Marcketta A., Mittelman D., Romanovitch M., Zhang C., Zheng-Bradley X., Abecasis G.R., McCarroll S.A., Flicek P., Underhill P.A., Coin L., Zerbino D.R., Yang F., Lee C., Clarke L., Auton A., Erlich Y., Handsaker R.E., Bustamante C.D. & Tyler-Smith C. 2016: Punctuated bursts in human male demography inferred from 1,244 worldwide Y-chromosome sequences. --- Nature Genetics 48: 593–599, doi:10.1038/ng.3559.

Rhie A., Nurk S., Cechova M., Hoyt S.J., Taylor D.J., Altemose N., Hook P.W., Koren S., Rautiainen M., Alexandrov I.A., Allen J., Asri M., Bzikadze A.V., Chen N.-C., Chin C.-S., Diekhans M., Flicek P., Formenti G., Fungtammasan A., Garcia Giron C., Garrison E., Gershman A., Gerton J.L., Grady P.G.S., Guarracino A., Haggerty L., Halabian R., Hansen N.F., Harris R., Hartley G.A., Harvey W.T., Haukness M., Heinz J., Hourlier T., Hubley R.M., Hunt S.E., Hwang S., Jain M., Kesharwani R.K., Lewis A.P., Li H., Logsdon G.A., Lucas J.K., Makalowski W., Markovic C., Martin F.J., Mc Cartney A.M., McCoy R.C., McDaniel J., McNulty B.M., Medvedev P., Mikheenko A., Munson K.M., Murphy T.D., Olsen H.E., Olson N.D., Paulin L.F., Porubsky D., Potapova T., Ryabov F., Salzberg S.L., Sauria M.E.G., Sedlazeck F.J., Shafin K., Shepelev V.A., Shumate A., Storer J.M., Surapaneni L., Taravella Oill A.M., Thibaud-Nissen F., Timp W., Tomaszkiewicz M., Vollger M.R., Walenz B.P., Watwood A.C., Weissensteiner M.H., Wenger A.M., Wilson M.A., Zarate S., Zhu Y., Zook J.M., Eichler E.E., O’Neill R.J., Schatz M.C., Miga K.H., Makova K.D. & Phillippy A.M. 2023: The complete sequence of a human Y chromosome. --- Nature 621: 344–354, doi:10.1038/s41586-023-06457-y.

Ross M.G., Russ C., Costello M., Hollinger A., Lennon N.J., Hegarty R., Nusbaum C. & Jaffe D.B. 2013: Characterizing and measuring bias in sequence data. --- Genome Biology 14: R51, doi:10.1186/gb-2013-14-5-r51.

Rosing-Asvid A., Löytynoja A., Momigliano P., Hansen R.G., Scharff-Olsen C.H., Valtonen M., Kammonen J., Dietz R., Rigét F.F., Ferguson S.H., Lydersen C., Kovacs K.M., Holland D.M., Jernvall J., Auvinen P. & Olsen M.T. 2023: An evolutionarily distinct ringed seal in the Ilulissat Icefjord. --- Molecular Ecology 32: 5932–5943, doi:10.1111/mec.17163.

Sundell T., Kammonen J., Mustanoja E., Biard V., Kunnasranta M., Niemi M., Nykänen M, Nyman T., Palo J.U., Valtonen M., Paulin L., Jernvall J. & Auvinen P. 2023: Genomic evidence uncovers inbreeding and supports translocations in rescuing the genetic diversity of a landlocked seal population. --- Conservation Genetics 24: 155–165, doi:10.1007/s10592-022-01497-9.

Tomaszkiewicz M., Medvedev P. & Makova K.D. 2017: Y and W chromosome assemblies: Approaches and discoveries. --- Trends in Genetics 33: 266–282, doi:10.1016/j.tig.2017.01.008.

Valtonen M., Palo J.U., Ruokonen M., Kunnasranta M. & Nyman T. 2012: Spatial and temporal variation in genetic diversity of an endangered freshwater seal. --- Conservation Genetics 13: 1231–1245, doi:10.1007/s10592-012-0367-5.

Valtonen M., Palo J.U., Aspi J., Ruokonen M., Kunnasranta M. & Nyman T. 2014: Causes and consequences of fine-scale population structure in a critically endangered freshwater seal. --- BMC Ecology 14: 22, doi:10.1186/1472-6785-14-22.

Wickham H. 2016: ggplot2: Elegant Graphics for Data Analysis. --- Springer-Verlag, New York.

Wutke S., Sandoval-Castellanos E., Benecke N., Döhle H.-J., Friederich S., Gonzalez J., Hofreiter M., Lõugas L., Magnell O., Malaspinas A.-S., Morales-Muñiz A., Orlando L., Reissmann M., Trinks A. & Ludwig A. 2018: Decline of genetic diversity in ancient domestic stallions in Europe. --- Science Advances 4: eaap9691, doi:10.1126/sciadv.aap9691.

Young A.C., Kirkness E.F. & Breen M. 2008: Tackling the characterization of canine chromosomal breakpoints with an integrated in-situ/in-silico approach: the canine PAR and PAB. --- Chromosome Research 16: 1193–1202, doi:10.1007/s10577-008-1268-9.

Yu G., Smith D.K., Zhu H., Guan Y. & Lam T.T.-Y. 2017: ggtree: an R package for visualization and annotation of phylogenetic trees with their covariates and other associated data. --- Methods in Ecology and Evolution 8: 28–36, doi:10.1111/2041-210X.12628.

