## Supplementary Information for "A chromosome-level reference genome assembly reveals extreme loss of Y-chromosomal diversity in the Saimaa ringed seal (*Pusa saimensis*)"

**Table S1.** Samples included in the analyses, with locality, sex, and year of collection. Regions are abbreviated as follows: PV = Pihlajavesi; KV = Kolovesi; HV = Haukivesi; SS = South Saimaa; NS = North Saimaa.

| Sample | Population | Region | Sex | Collection year |
|---|---|---|---|---|
| 1090 | Saimaa | PV | F | 1986 |
| 1235 | Saimaa | SS | M | 1987 |
| 1262 | Saimaa | HV | M | 1988 |
| 1270 | Saimaa | HV | F | 1988 |
| 1271 | Saimaa | PV | M | 1988 |
| 1482 | Saimaa | KV | F | 1994 |

| | | | | |
|---|---|---|---|---|
| 1488 | Saimaa | SS | M | 1994 |
| 1552 | Saimaa | PV | F | 1995 |
| 1957 | Saimaa | PV | F | 1997 |
| 1977 | Saimaa | PV | F | 1997 |
| 1983 | Saimaa | HV | M | 1997 |
| 2232 | Saimaa | HV | M | 2000 |
| 2233 | Saimaa | PV | M | 2000 |
| 2259 | Saimaa | SS | F | 2001 |
| 2277 | Saimaa | HV | F | 2001 |
| 2332 | Saimaa | SS | F | 2003 |
| 2341 | Saimaa | HV | F | 2003 |
| 2342 | Saimaa | HV | M | 2003 |
| 2343 | Saimaa | PV | F | 2003 |
| 2348 | Saimaa | KV | M | 2003 |
| 2350 | Saimaa | HV | F | 2003 |
| 2357 | Saimaa | SS | M | 2004 |
| 2379 | Saimaa | HV | F | 2005 |
| 2382 | Saimaa | HV | M | 2005 |
| 2392 | Saimaa | HV | F | 2005 |
| 2395 | Saimaa | PV | F | 2006 |
| 2396 | Saimaa | HV | M | 2006 |
| 2397 | Saimaa | PV | F | 2006 |
| 2399 | Saimaa | KV | F | 2006 |
| 2405 | Saimaa | PV | M | 2006 |
| 2427 | Saimaa | PV | M | 2007 |
| 2432 | Saimaa | HV | M | 2007 |
| 2433 | Saimaa | HV | F | 2007 |
| 2436 | Saimaa | SS | M | 2007 |
| 2452 | Saimaa | HV | M | 2008 |
| 2474 | Saimaa | NS | M | 2009 |
| 2496 | Saimaa | PV | F | 2009 |
| 2499 | Saimaa | HV | M | 2010 |
| 2506 | Saimaa | HV | M | 2010 |

| | | | | |
|---|---|---|---|---|
| 2509 | Saimaa | PV | F | 2010 |
| 2511 | Saimaa | HV | M | 2010 |
| 2513 | Saimaa | PV | M | 2010 |
| 2514 | Saimaa | PV | M | 2011 |
| 2528 | Saimaa | PV | F | 2011 |
| 2550 | Saimaa | PV | M | 2012 |
| 2551 | Saimaa | HV | F | 2012 |
| 2557 | Saimaa | HV | M | 2012 |
| 2564 | Saimaa | PV | M | 2013 |
| 2565 | Saimaa | PV | M | 2013 |
| 2566 | Saimaa | SS | M | 2013 |
| 2567 | Saimaa | PV | M | 2013 |
| 2568 | Saimaa | HV | F | 2013 |
| 2569 | Saimaa | PV | M | 2013 |
| 2570 | Saimaa | PV | M | 2013 |
| 2574 | Saimaa | PV | M | 2013 |
| 258 | Saimaa | NS | M | 1980 |
| 2585 | Saimaa | HV | F | 2013 |
| 2614 | Saimaa | PV | F | 2014 |
| 292 | Saimaa | KV | M | 1981 |
| 297 | Saimaa | NS | M | 1981 |
| 298 | Saimaa | HV | M | 1981 |
| 819 | Saimaa | NS | F | 1985 |
| BG-02 | Baltic | Bothnian Bay | M | 2013 |
| BG-03 | Baltic | Bothnian Bay | M | 2013 |
| L1683 | Ladoga | Karvatsusaari | M | 1996 |
| LG-01 | Ladoga | Vidlitsa | M | 2014 |
| LG-02 | Ladoga | Vidlitsa | M | 2015 |
| N3-09 | Baltic | Bothnian Bay | M | 2009 |
| N7-07 | Baltic | Bothnian Bay | M | 2007 |
| NN3-07 | Baltic | Bothnian Bay | M | 2007 |
| PH027 | Greenland | Qaanaaq | M | NA |
| PH043 | Greenland | Ittoqqortoormiit | M | NA |

| | | | | |
|---|---|---|---|---|
| PH046 | Greenland | Qaanaaq | M | NA |
| PH047 | Greenland | Ittoqqortoormiit | M | NA |
| PH063 | Greenland | Qaanaaq | M | NA |
| PH082 | Greenland | Ittoqqortoormiit | M | NA |
| PH085 | Greenland | Ittoqqortoormiit | M | NA |
| S11-07 | Svalbard | Tempelfj. | M | 2007 |
| S20-07 | Svalbard | Tempelfj. | M | 2007 |
| S4-07 | Svalbard | NA | NA | NA |

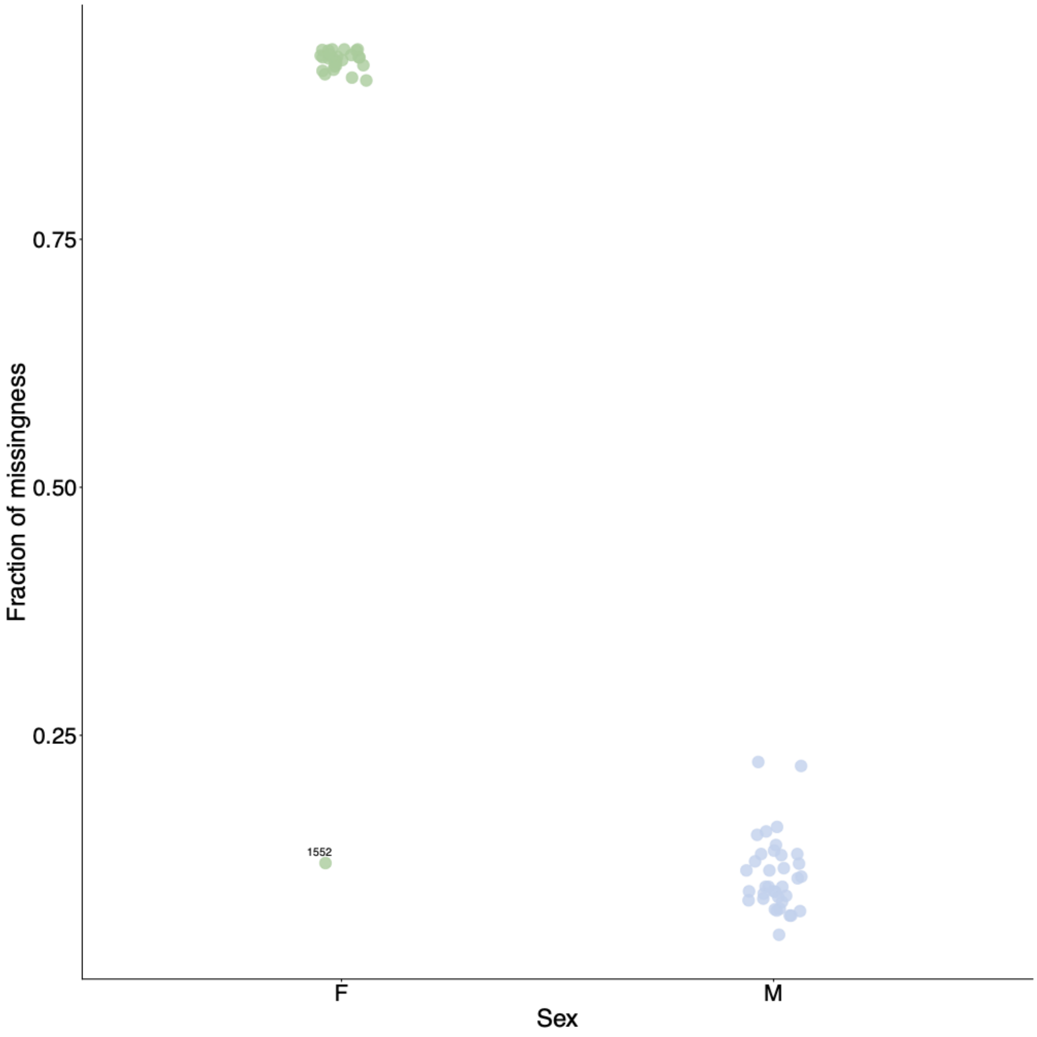


**Fig. S1.** Fraction of missing Y-chromosomal variants in 25 female and 37 male Saimaa ringed seals, showing a clear separation between the sexes based on this metric. Individual 1552, a stillborn pup potentially misassigned as female, is indicated.

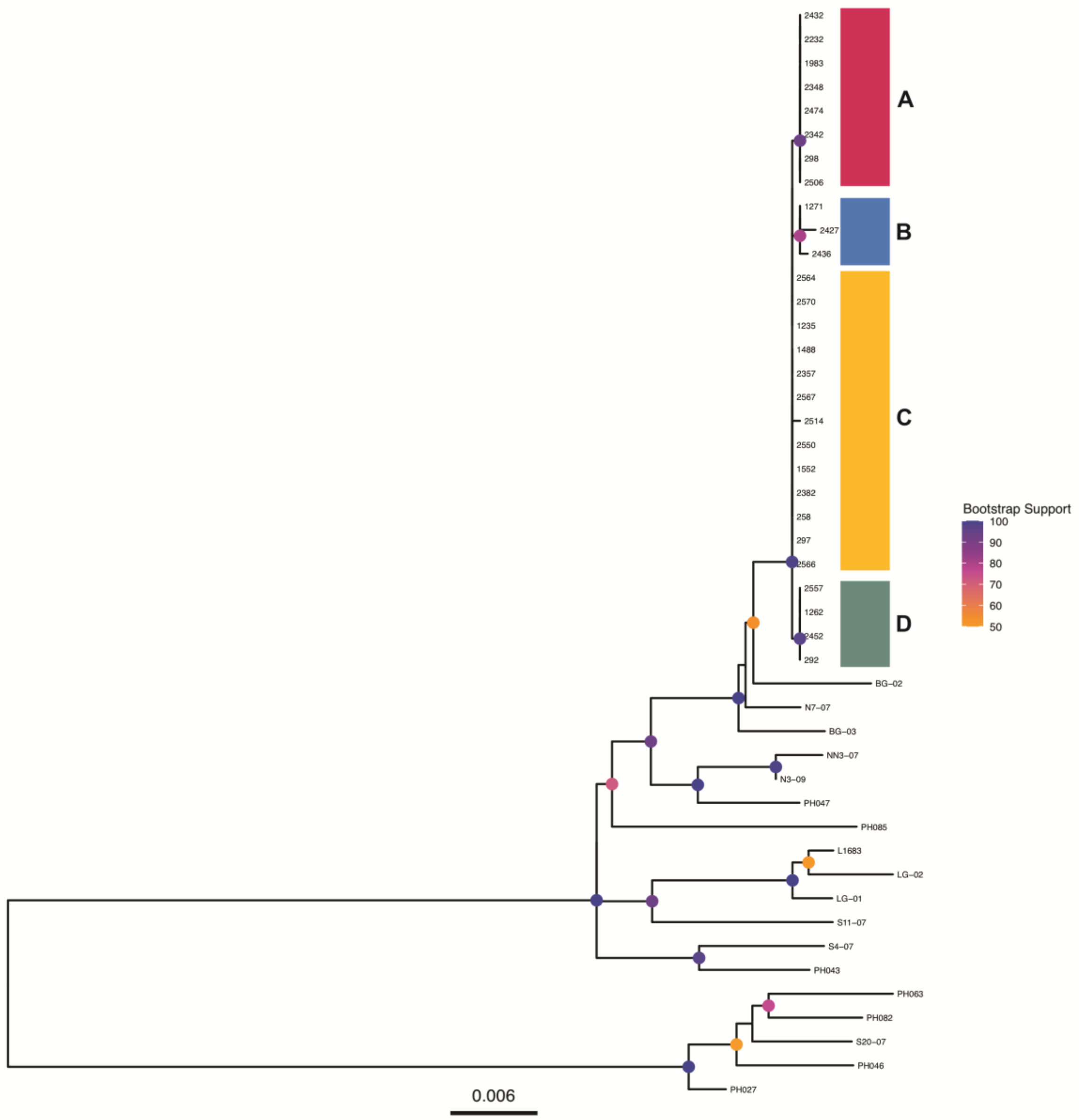


**Fig. S2.** Y-chromosomal phylogenetic tree of 46 male ringed seals, with missing genotypes assigned as reference calls, showing bootstrap support values. Saimaa ringed seals form a monophyletic clade comprising four major Y chromosome haplotypes, after haplotypes represented by single individuals were collapsed to their ancestral haplotype groups. Haplotype assignment and colour-coding as in Fig. 3.

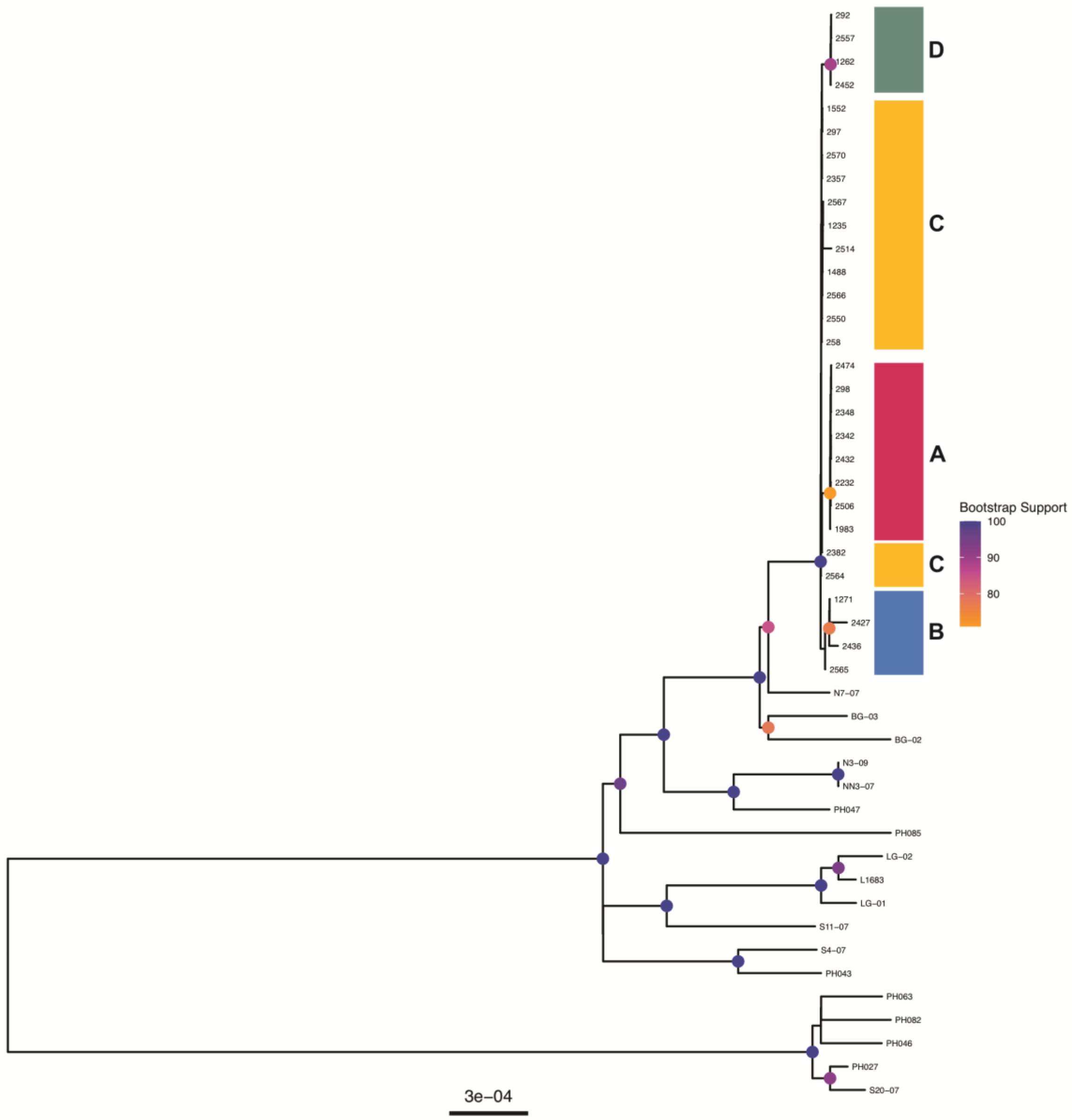


**Fig. S3.** Y-chromosomal phylogenetic tree of 46 male ringed seals, with missing genotypes retained as "N", showing bootstrap support values. Four main Saimaa clades are visible similarly to the Fig. S2.

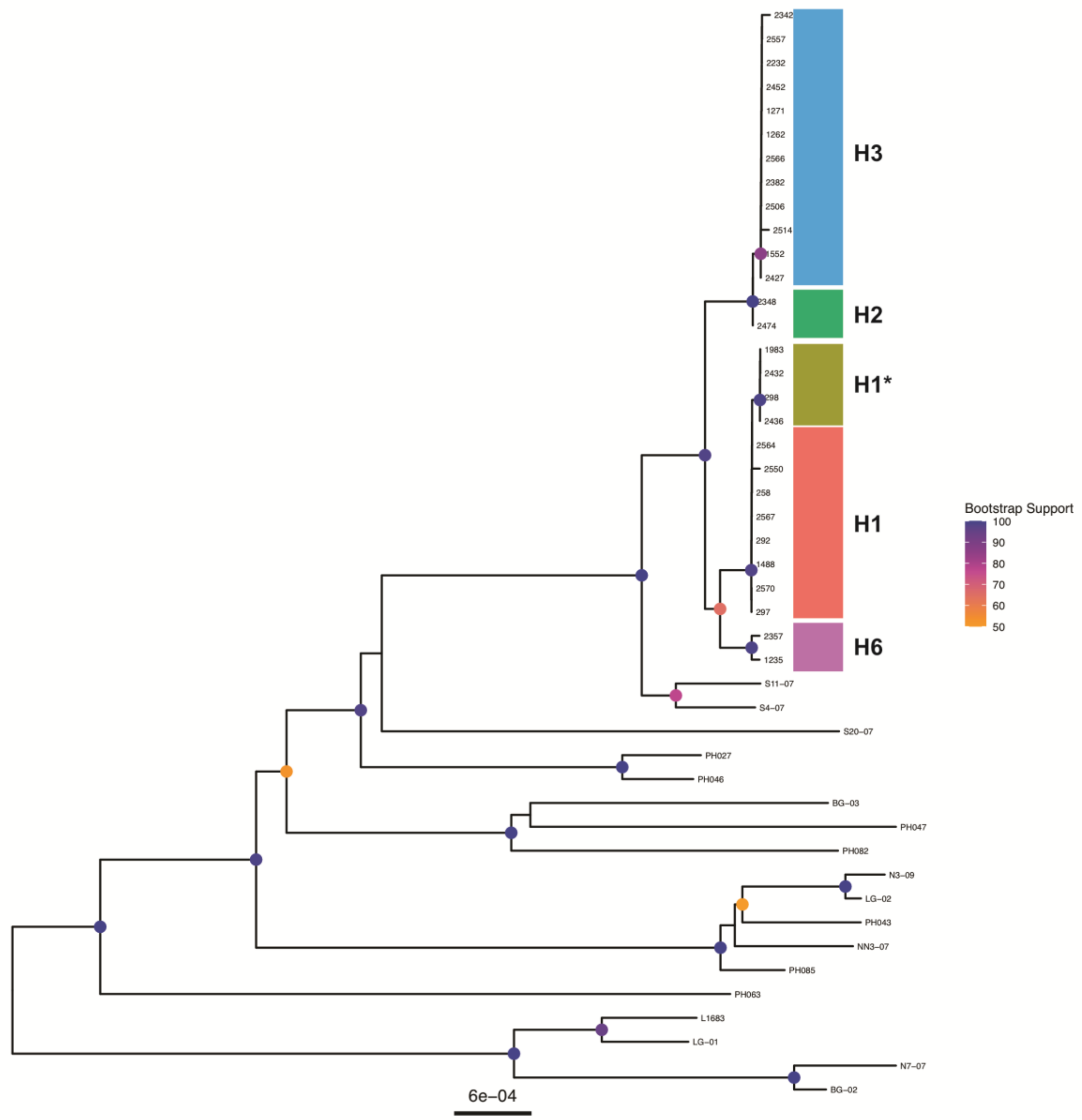


**Fig. S4.** Mitochondrial phylogenetic tree of 46 male ringed seals, showing bootstrap support values. Saimaa ringed seals form a monophyletic clade comprising five major mitochondrial haplotypes, after haplotypes represented by single individuals were collapsed to their ancestral haplotype groups.
